# Mapping of Quantitative Trait Loci for Traits linked to Fusarium Head Blight Symptoms Evaluation In Barley RILs

**DOI:** 10.1101/751552

**Authors:** Piotr Ogrodowicz, Anetta Kuczyńska, Krzysztof Mikołajczak, Tadeusz Adamski, Maria Surma, Paweł Krajewski, Hanna Ćwiek-Kupczyńska, Michał Kempa, Michał Rokicki, Dorota Jasińska

## Abstract

Fusarium head blight (FHB) is a devastating disease in small grain cereals worldwide. The disease results in the reduction of grain yield and affects its quality. In addition, mycotoxins accumulated in grain are harmful to both humans and animals. It has been reported that response to pathogen infection may be associated with the morphological and developmental characteristics of the host plant, e.g. the earliness and plant height. Despite the many studies the effective markers for the selection of barley genotypes with increased resistance to FHB have not thus far been developed. Therefore, exploring the genetic relationship between agronomic traits (e.g. heading date or stem height) and disease resistance is of importance to the understanding of plant resistance via “diesease escape” or dwarf stature. The studied plant material consisted of 100 recombinant inbred lines (RIL) of spring barley. Plants were examined in field conditions (three locations) in a completely randomized design with three replications. Barley genotypes were artificially infected with spores of *Fusarium* before heading. Apart from the main phenotypic traits (plant height, spike characteristic, grain yield) the infected kernels were visually scored and the content of deoxynivalenol (DON) mycotoxin was investigated. A set of 70 Quantitative Trait Loci (QTLs) were detected through phenotyping of the mapping population in field condition and genotyping using a barley Ilumina iSelect platform with 9K markers. Six loci were detected for FHB index on chromosomes 2H, 3H, 5H and 7H. The region on the short arm of the 2H chromosome was detected in the current study, in which many QTLs associated with FHB- and yield-related characters were found. This study confirms that agromorphological traits are tightly related to the FHB and should be taken into consideration when breeding barley plants for FHB resistance.

## Introduction

Fusarium head blight (FHB) or a scabs affects different species of crops around the world. The infection is caused by several fungal pathogens, among others *Fusarium culmorum* (W. G. Sm.) *Sacc* and *Fusarium graminearum* (teleomorph stage: *Gibberella zeae*). The first species of *Fusarium* has been found to dominate in regions with warm and humid conditions, whereas the second has been associated with cool, wet and humid conditions [1]. The visible symptoms of the disease are bleaching of some of the florets in the head before maturity stage. Other symptoms include tan to brown discoloration at the base of the spike and a pink or orange colored mold at the base of the florets under moist conditions. Kernels observed on the infected spikes are shriveled, white, and chalky in appearance. Moreover, *Fusarium* spp. produce trichothecene - deoxynivalenol (DON) [2]. This mycotoxin disrupts normal cell function by inhibiting protein synthesis [3] which can result in reducing grain quality and yield performance. Floret sterility and deformed kernels contribute to significant yield loss [4]. In Europe 15 – 55% of the barley products are contaminated with DON [5].

DON poses a real threat to human and livestock health. This mycotoxin is also known as “vomitoxin” due to its emetic effects after consumption [6]. DON levels present in barley (*Hordeum vulgare* L.) and wheat (*Triticum aestivum* L.) infected with FHB may vary according to the time of infection and environmental factors. It is well known that infection is favored by moist and warm conditions [7, 8]. While the presence of scab can be determined through visual inspection, the presence of DON cannot. The assessment of disease severity is based on the ratio of symptomatic spikelets on each spike and the proportion of infected spikes among the tested plants [9]. Although this method is widely used in the screening of resistant germplasms, the results are subjective. Hence, different types of chromatography for identification and quantification of mycotoxins in barley are commonly in use [10, 11]. However, due to the time-consuming and costly nature of these methods, commercial immunometric assays, such as enzyme-linked immunosorbent assay (ELISA), are frequently used for the monitoring of mycotoxin content [12, 13].

Disease control is achieved by the deployment of resistant cultivars. However, breeding for FHB resistance has proved difficult due to the complex inheritance of the resistance genes [14] and the strong genotype-by-environment interaction [15].

One of the several crop species most vulnerable to FHB infection is barley (*Hordeum vulgare* L.). This species is a cereal crop of major importance, ranked the fourth grain crop in the world in terms of production volume [16]. Its major uses are both as animal feed and as a component of human nutrition [17, 18]. In addition, barley is perceived to be a model plant in genetic study due to genome colinearity and synteny across rye, barley and wheat [19].

Fusarium poses a real threat for barley plants especially in regions that are prone to have long periods of wet weather during the flowering stage [4]. Host plants are most vulnerable to infection during anthesis due to development of fungal spores on anthers and polen containg nutrients [20]. Numerous morphological traits have been shown to be associated with FHB resistance in barley [21]. Heading date, plant height, and spike characters (linked to spike compactness) are mostly investigated [22, 23]. Days to heading is often negatively correlated with FHB susceptibility and usually results in disease escape [24]. Hence, using the least susceptible varieties with different flowering date may reduce FHB risks. Two categories of resistance to FHB are generally recognized: type I (resistance to initial infection) and type II (resistance to fungal spread within the spike) [25]. Another kind of resistance has been described as a third type and is related to the accumulation of mycotoxins within the grains [26].

Studies designed to determine the number and chromosomal location of loci contributing to FHB resistance and the accumulation of DON are urgently needed for the resistance breeding efforts. Resistance to FHB is a complex trait controlled by multiple genes and affected by environmental factors [27, 28]. QTL have been identified for both qualitative and quantitative disease resistance in wheat and barley [4]. Resistance to FHB and DON level content have been mapped to all seven barley chromosomes [29, 30]. The most common regions related to FHB resistane have been previously reported on chromosomes 2H and 6H in many studies [3, 25, 31]. Other traits including awned/awnless ears [26] and spike compactness [32] have also been studied. Plant height is another parameter frequently investigated, and a negative correlation of this trait with Type I FHB susceptibility has been frequently documented [33].

Molecular markers have become increasingly important for plant genome analysis. Different classes of DNA markers have been developed and implemented over time [34]. A new genotyping platform followed in 2009 that introduced larger numbers of markers based on SNP discovery in Next Generation Sequencing data when the Illumina’s oligo pool assay as a marker platform [35] was designed to improve the genotyping process. The 9K iSelect chip contains 7864 SNPs [36] and enables higher efficiency and cost reduction. In the current study this chip was employed due to the favorable tradeoff between genotyping costs and marker density.

This study aimed to map quantitative traits loci linked to agronomic properties in mapping population grown in field conditions and subjected to artificial *Fusarium* infection. Evaluation of diease severity was based on both visual assessment of infection and evaluation of deoxynivalenol content.

## Material and methods

### Plant material

A 100-RIL population of spring barley obtained from the cross between the Polish cultivar Lubuski and a Syrian breeding line (Cam/B1/CI08887//CI05761) was studied in field conditions, together with both parental forms. The plant materials were described in detail in Ogrodowicz et al. [37].

### Field experiment

The experiments were perfomed in experimental areas belonging to Poznan Plant Breeding Company (PPB) in three locations: Nagradowice (NAD –Western Poland, 52°19′14″N, 17°08′54″E), Tulce (TUL - Western Poland, 52°20′35.2″N 17°04′32.8″E), Leszno (LES - Western Poland, 51°50′45″N 16°34′50″E). At each location, the experiments were performed in randomized blocks with three replications. The effects of the *Fusarium* infection were evaluated during the 2016 growing season. The two experimental variants were: V1 – variant 1 – control condition, V2 – variant 2 – inoculation. Control rows were established at a distance of 20.0 m from the plots designated for inoculation. This isolation was necessary to protect the plants against infection during inoculation.

### Methodology

Inoculum was prepared just before the inoculations by liquid cultures of *Fusarium culmorum* (isolate KF846) and 0.0125% of TWEEN®20 (Sigma-Aldrich Chemie GmbH). Conidia concentration was adjusted to 10^5^/1 mL. Inoculation was performed at flowering stage (BBCH scale 61). After inoculation the plants were micro-irrigated for three days to maintain moisture.

### Agronomic traits

At maturity, the number of spikelets (NSS), number of kernels (NGS), length of spike without awns (LS) and grain weight per spike (GWS) were observed on 10 randomly selected plants. In addition, during investigation, heading date (HD), plant height (HD) and stature of plants (Stature) were recorded. Finally, the plots were harvested and grains from trials were weighed to derive grain yield per plot (GY). In our study, two additional traits were added to the analysis: the numbers of sterile spikelets per spike (Sterility) and spike density (Density). The measured traits with ontology annotation are listed in Table 1.

**Table 1.** List of phenotypic traits with description, abbreviations, measured units and ontology annotation.

| Trait (unit) | Trait description | Abbrev | Annotation |
| --- | --- | --- | --- |
| Number of spikelets per spike | Number of spikelets in spike from 10 randomly selected spikes in a pot | NSS | http://purl.obolibrary.org/obo/TO_0000456 |
| Number of grains per spike | Number of grains collected from 10 randomly selected spikes in a pot | NGS | http://purl.obolibrary.org/obo/TO_0002759 |
| Length of spike (cm) | Length of spike from 10 randomly selected spikes in a pot (without awns) | LS | http://purl.obolibrary.org/obo/TO_0000040 |
| Fertile and sterile spikelets number per spike | Fraction of sterile spikelets per spike, calculated as a ratio of number of spikelets per spike (NSS) to number of grains per spike (NGS) | Sterility | http://purl.obolibrary.org/obo/TO_0000436 |
| Spike density/the number of spikelets per unit length (centimeter) of spike | Trait calculated by dividing the number of spikelets per spike by the length of the spike | Density | http://purl.obolibrary.org/obo/TO_0020001 |
| Grain weight per spike (g) | Average weight of grain per spike, calculated from 10 randomly selected spikes in a pot | GWS | http://purl.obolibrary.org/obo/TO_0000589 |
| Grain yield (g) | Weight of grain harvested per plot | GY | http://purl.obolibrary.org/obo/TO_0000396 |
| 1000-grain weight (g) | Average weight of 1000 grains, calculated as 1000 * average weight of one grain for spikes in a pot | TGW | http://purl.obolibrary.org/obo/TO_0000382 |
| Heading date (number of days) | Number of days from sowing to emergence of inflorescence (spike) from the flag leaf (BBCH), assessed when spikes emerged on at least 50% of plants | HD | http://purl.obolibrary.org/obo/TO_0000137 |
| Length of main stem (cm) | Average of measurements of length of stem from ground level to the end of spike (without awns) for 10 randomly selected plants in a pot | LSt | http://purl.obolibrary.org/obo/TO_0000576 |
| Stature | Overall size/ shape of plant | Stature |  |
| FHB index | Spike infection in % percentage of spikelets affected within a spike / percentage of infected spikes per plot)*100 | FHBi | http://purl.obolibrary.org/obo/TO_0000662 |
| DON concentration (ppb) | Deoxynivalenol content of the grain | DON | http://purl.obolibrary.org/obo/TO_0000669 |
| Number of damaged kernels | Number of kernels classified as damaged (pinkish or discoloured) per 10 randomly selected spikes | FDKn |  |
| Weight of damaged kernels (g) | Weight of kernels classified as damaged (pinkish or discoloured) per 10 randomly selected spikes | FDKw |  |
| Number of healthy kernels | Number of kernels classified as healthy per 10 randomly selected spikes | HLKn |  |
| Weight of healthy kernels (g) | Weight of kernels classified as healthy per 10 randomly selected spikes | HLKw |  |

### Disease symptoms evaluation

Disease development was visually scored using the Fusarium Head Blight index (FHBi) (percentage of infected spikelets within a spike / percentage of infected spikes per plot) ×100. After harvest Fusarium-damaged kernels (FDK) were observed - the number (FDKn) and weight (FDKw) of kernels, which were classified as pinkish or discoloured (Fig 1, 2). Those kernels that appeared to be healthy were scored as HLK (healthy looking kernels – division: HLKn and HLKw). FDK and HLK rate was estimated for infected and controlled kernels in one location (NAD).

**Fig 1.**
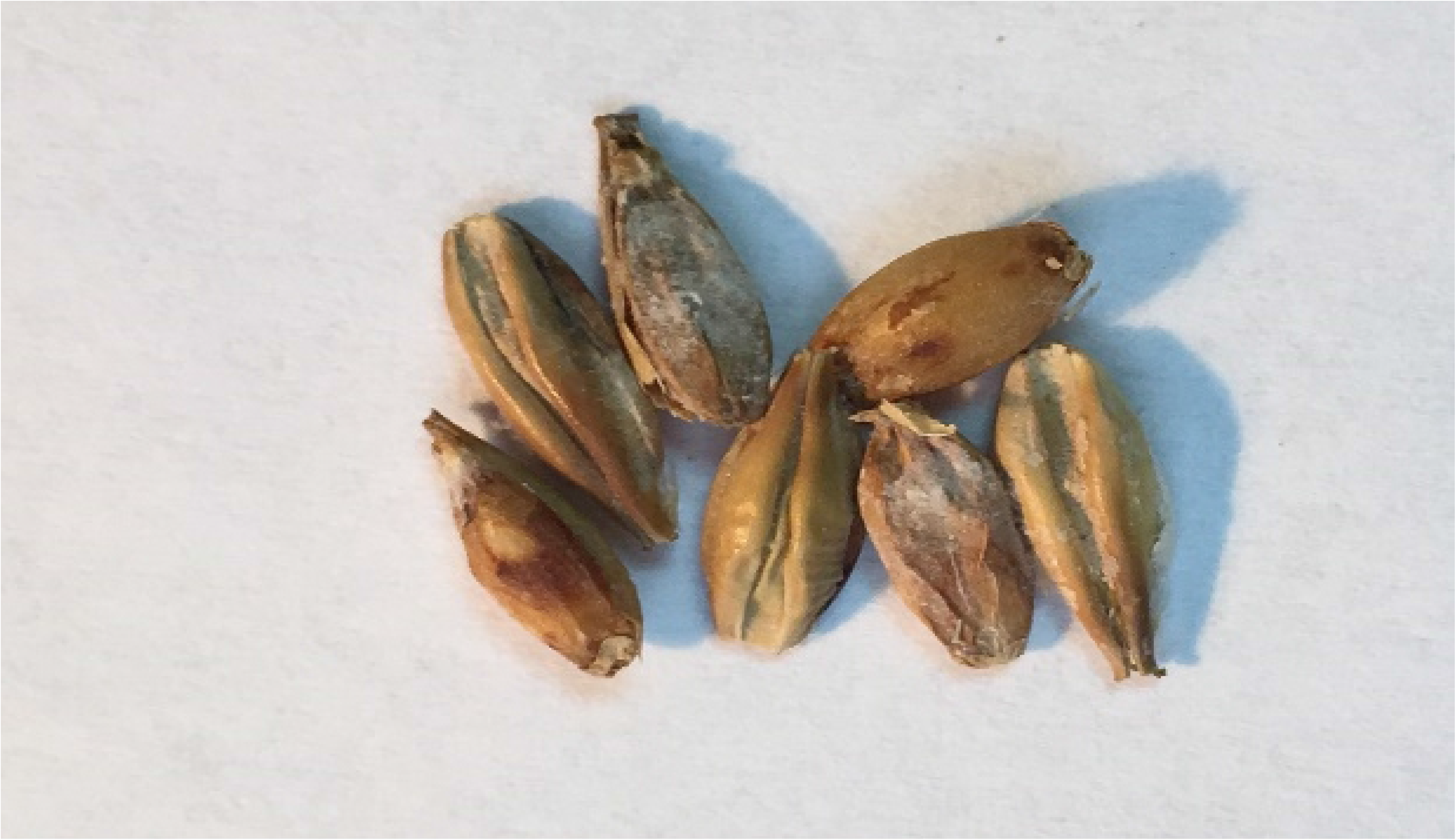
Seeds, observed in Lcam plants, with moderate or severe *Fusarium* symptoms. Seeds are thin, with some dark discolouration. This image was captured at 40 x magnification under the Motic BA410-E microscope.

**Fig 2.**
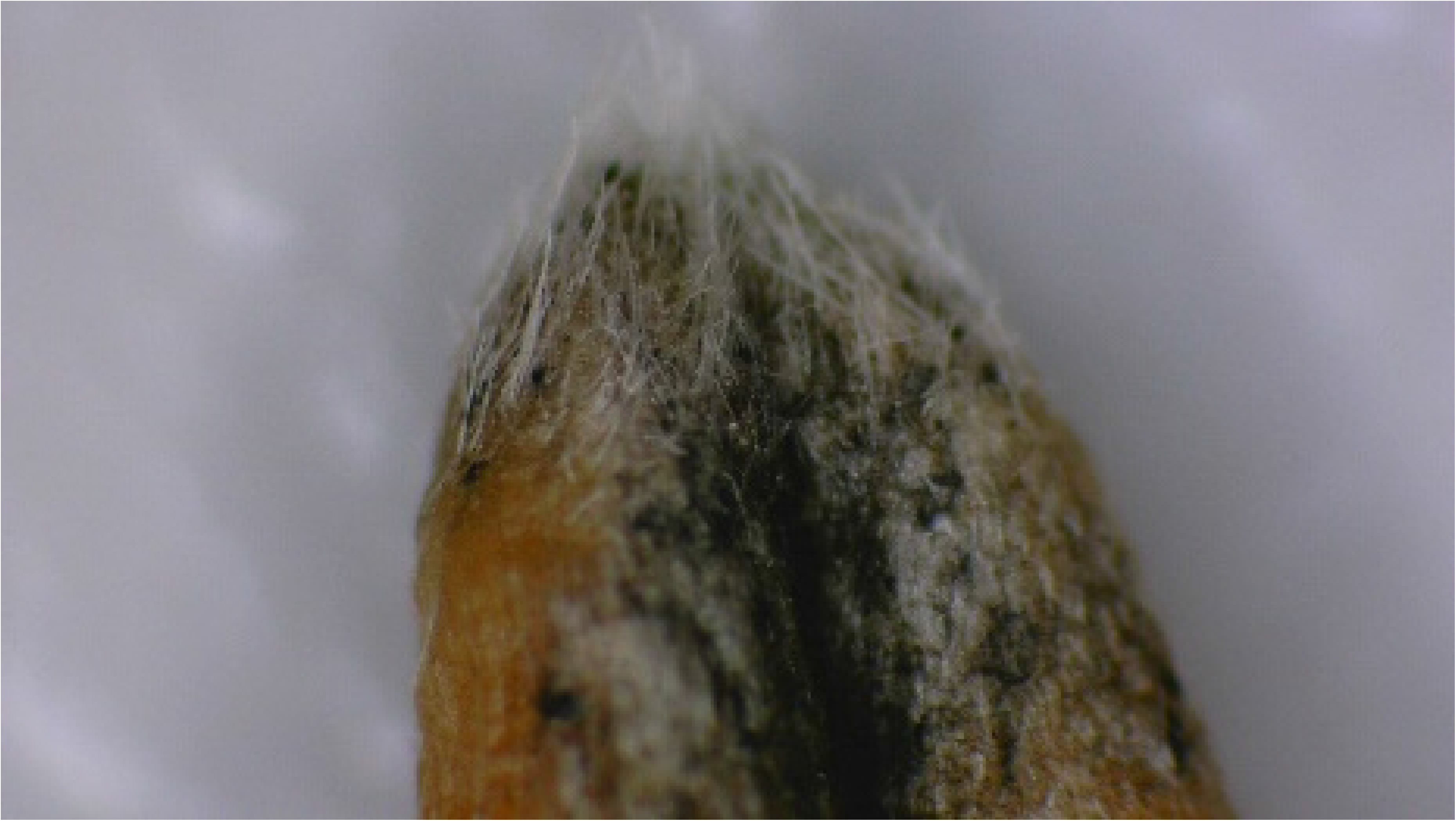
Abundant mycelial growth observed on the grain surface. This image was captured at 40 x magnification under the Motic BA410-E microscope.

DON content (mg kg^−1^ ppm) from infected grain samples (each experiment with three replications) was assessed using a Ridascreen®DON competitive enzyme immunoassay kit (R-Biopharm AG, Darmstadt, Germany) according to the manufacturer’s instructions. Absorbance was measured at 450 nm with a spectrophotometer (Chromate Microplate Reader). The data were evaluated with RIDA®SOFT Win software. Within a single locations (NAD, TUL, LES) samples obtained from plants grown in controlled conditions (exposed to natural infection) were pooled together into one repetition and this pool was assayed as above.

### Genotyping

Genomic DNA was extracted from young leaf tissue as described in Mikołajczak et al. 2016 [38]. DNA quantity and concentration were measured with a NanoDrop 2000 (Thermo Scientific™). The DNA samples were diluted to ∼ 50 ng/µL and sent to Trait Genetics, Gatersleben, Germany (http://www.traitgenetics.com). The barley iSelect SNP chip contains a total of 7.842 SNPs that comprise 2.832 of the existing barley oligonucleotide pooled assay (BOPA1 and BOPA2) SNPs discovered and mapped previously [39, 40], plus 5.010 new SNPs developed from Next Generation Sequencing data [36, 41]. SNPs which were not polymorphic between the parents, contained more than 10% of missing values, or with minor allele frequency smaller than 15% were removed from the markers set.

### Map construction

Genetic maps were calculated using the JoinMap 4.1 software [42]. All markers were analyzed for their goodness of fit using a chi-square test at significance level α = 0.05. A segregation ratio of 1:1 was expected. Markers with other segregation ratios were categorized as odd. The markers which were mapped to incorrect regions of the chromosomes were removed from the mapping and the marker order was calculated again. The localization of markers was designated using the maximum likelihood algorithm command. Markers were assigned to linkage groups applying the independence LOD (logarithm of the odds) parameter with LOD threshold values ranging from 6.0 to 9.0. The recombination frequency threshold was set at level ≤4. Recombination fractions were converted to map distances in centimorgans (cM) using the Kosambi mapping function. A map was drawn using MapChart 2.2

### Data analysis and QTLs mapping

Observations for RILs were processed by analysis of variance in a mixed model with fixed effects for location, treatment and location × treatment interaction, and with random effects for line and interaction of line with location and interaction of line with location and treatment. The residual maximum likelihood algorithm was used to estimate variance components for random effects and the F-statistic was computed to assess the significance of the fixed effects. Pearson correlation coefficients between all the analyzed traits were calculated. QTL analysis was performed for the linkage map with the mixed model approach described by Malosetti et al. [43], including optimal genetic correlation structure selection and significance threshold estimation. The threshold forthe−log10(P-value) statistic was computed by the method of Li and Ji [44] to ensure the genome-wide error rate was less than 0.01. Selection of the set of QTL effects in the final model was performed at P<0.05; the P-values for the Wald test were computed as the mean from the values obtained by adding and dropping the QTL main and interaction effects in the model. All the above computations were performed in Genstat 16 [45]. RIL lines where the lack of genotypic data did not exceed 20/15% were used to map QTL. QTL identification was performed for all studied traits.

The detected QTLs were labeled using a system described for wheat and *Arabidopsis* [46, 47], with minor modifications. The QTLs names consist of the prefix Q followed by a two- or three-letter descriptor of the phenotype (abbreviation of the trait name), an indicator for the laboratory, the number of the chromosome and a serial number. For traits linked to FDK and HLK the QTL names were extended by adding the letter “w” or “n” for loci found for trait weight of FDK, HLK and number of FDK, HLK, respectively.

QTL effects in individual trials were considered major if the fraction of explained variance exceeded 12.32% (upper quartile of the distribution of explained variance) according to the rules employed by [48] and [49] (with minor modifications).

The barleymap pipeline (http://floresta.eead.csic.es/barleymap) [50] was used to identify potential candidate genes underlying the particularly robust QTL of this study. Markers from seven regions harboring QTLs for studied traits were annotated and the gene search was extended to an interval of ±2cM around markers. Overrepresentation analysis (ORA) of GO terms (among high confidence gene class) was performed using the hypergeometric distribution [51, 52] and applying the Benjamini–Hochberg correction (FDR level of 0.05) [53].

## Results

### Phenotypic analysis

The parents of the LCam population were characterized with 11 agronomical traits under two different conditions (infection and control treatments). Evaluation of disease severity was studied by using measurements of six FHB-related traits in both type of previously mentioned conditions. The distributions of trait values among RILs are visualized in Fig 3.

**Fig 3.**
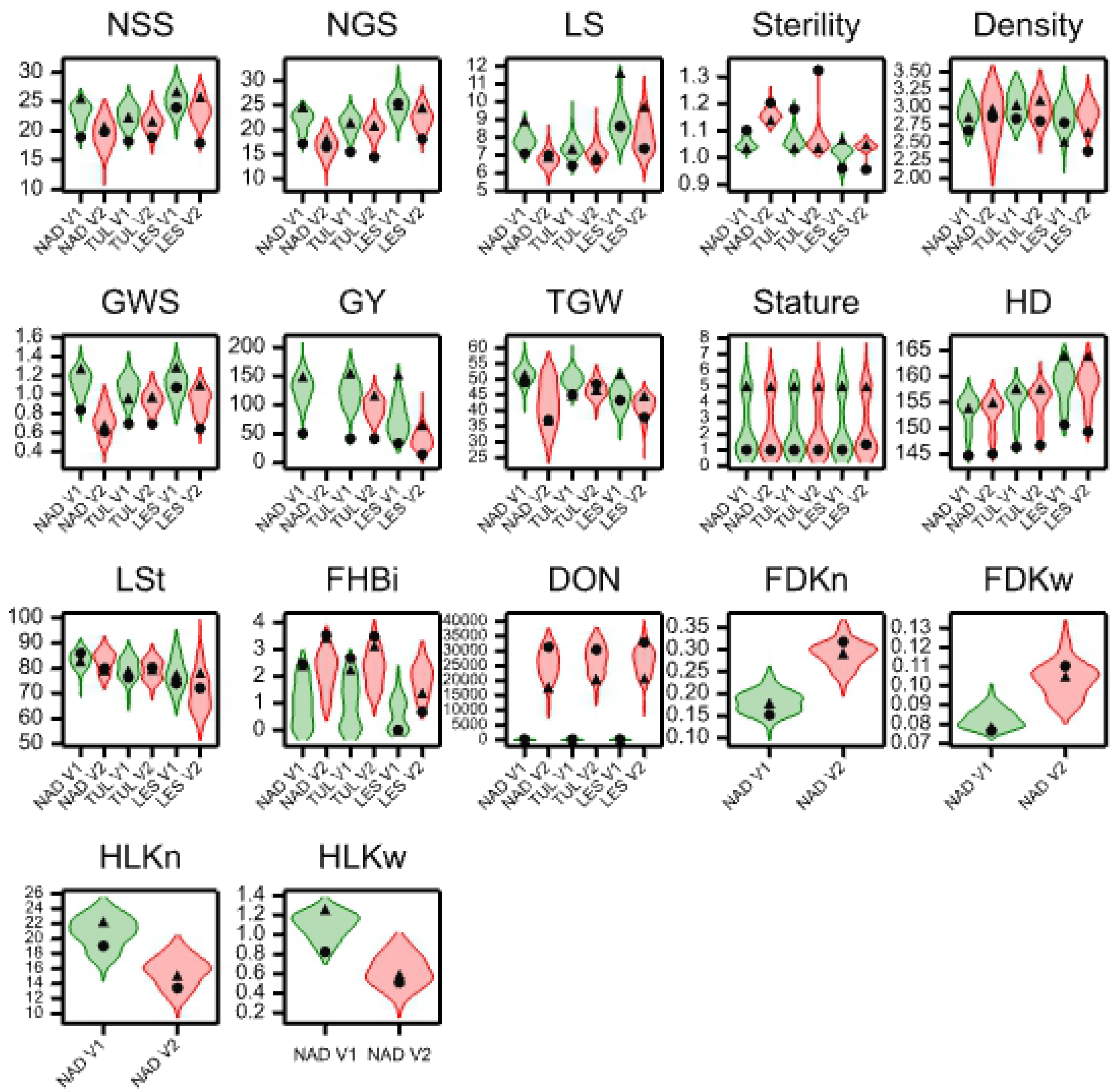
Violin plots for studied traits measured in the LCam population in control (V1, green color) and infected (V2, red color) conditions in three locations. Black symbols: triangle - Lubuski, dot - CamB.

The parental forms were differentiated in terms of all the studied characters (S1 Table). Lubuski showed higher mean values of traits linked to yield performance (e.g. GWS, GY,). The Syrian genotype showed a lower mean value of HD in all trials and under both types of treatments (heading for CamB was 11 days earlier than for Lubuski).

A substantial GY decline was observed for Lubuski in infection condition (40.1%). In comparison, for CamB a lower relative decline for GY was observed (17.3%). Both mean value for FHBi and traits associated with visual evaluation of *Fusarium* symptoms (FDK) increased in the infection conditions. For the Syrian parent a higher mean value of DON concentration was noted in comparison to that of European parent. For both parental forms low concentrations of mycotoxin were also observed in control conditions.

The mean values of the studied traits for RILs are presented in S2 Table. Relatively high values of variation coefficients were observed in location NAD under infection for traits: NSS, NGS, Density, GWS and TGW. In the LES location very high values of CV were noted for traits FHBi and DON in control conditions.

FHBi varied across locations with the mean FHBi ranging from 1.89 to 2.26 under infection treatment and from 0.62 to 0.99 in control condition (S2 Table). The amount of DON, measured in grains from infected plants, varied from a maximum of 39990.00 μg kg−1 (TUL) to 8060.00 μg kg−1 (NAD). Mean DON values of 26 439.47, 25 684.27 and 27 144.47 μg kg−1 for infection treatment in LES, NAD, TUL were observed, respectively. In control conditions relatively high coefficients of variation were noted for DON and FHBi.

Analysis of variance indicated significant effects of location and treatment on the RIL population for all traits (P<0.001) with several exceptions (Table 2). In all cases, the variance components for all types of interactions were smaller than those for lines. For FHBi a significant line × location interaction was noted. No signicant interaction was observed for line × treatment in this case. An insigificant effect was noticed in terms of the interaction line × location for DON content.

**Table 2.** ANOVA results and variance components estimated for studied traits.

| Trait (abbrev. | P-values for effects of |  |  | Variance components and std. errors for |  |  |  |  |  |  |  |
| --- | --- | --- | --- | --- | --- | --- | --- | --- | --- | --- | --- |
|  | location | treatment | location ×<br>treatment<br>interacton | lines | s.e. | interaction line x<br>location | s.e. | interaction line x<br>treatment | s.e. | interaction line<br>location × treatm | s.e. |
| NSS | <0.001 | <0.001 | <0.001 | 2.447 | 0.397 | 0.32 | 0.135 | 0.02 | 0.093 | 0 | - |
| NGS | <0.001 | <0.001 | <0.001 | 2.766 | 0.446 | 0.394 | 0.143 | 0.033 | 0.096 | 0 | - |
| LS | <0.001 | <0.001 | <0.001 | 0.2643 | 0.0443 | 0.0634 | 0.0178 | 0 | - | 0.0181 | 0.0194 |
| Sterility | <0.001 | <0.001 | <0.001 | 0.000139 | 0.00005 | 0.000251 | 0.000065 | 0 | - | 0 | - |
| Density | <0.001 | 0.138 | 0.009 | 0.0256 | 0.00448 | 0.0056 | 0.00176 | 0.0022 | 0.00135 | 0 | - |
| GWS | <0.001 | <0.001 | <0.001 | 0.00969 | 0.00167 | 0.00144 | 0.0065 | 0.00065 | 0.00052 | 0 | - |
| GY | <0.001 | <0.001 | 0.006 | 329.9 | 52.8 | 0 | - | 0 | - | 0 | - |
| TGW | <0.001 | <0.001 | <0.001 | 1.9 | 0.87 | 1.22 | 1.03 | 0.97 | 0.88 | 0 | - |
| Stature | 0.927 | 0.15 | 0.31 | 3.05756 | 0.44322 | 0.25067 | 0.02747 | 0 | - | 0.02229 | 0.00496 |
| HD | <0.001 | <0.001 | 0.035 | 9.625 | 1.419 | 1.142 | 0.134 | 0.013 | 0.024 | 0.019 | 0.041 |
| LSt | <0.001 | <0.001 | <0.001 | 9.55 | 2.047 | 5.118 | 1.414 | 0.519 | 0.881 | 11.234 | 1.425 |
| FHBi | <0.001 | <0.001 | 0.02 | 0.0471 | 0.0325 | 0.4351 | 0.0461 | 0.0045 | 0.0037 | 0.0187 | 0.0056 |
| DON<sub>x</sub> | <0.001 |  |  | 21271008 | 3204801 | 0 | - | - | - |  |  |
| FDKn | - | <0.001 | - | 0.000228 | 0.000082 | - | - | 0.000189 | 0.000084 | - | - |
| FDKw | - | <0.001 | - | 0.0000157 | 0.00000595 | - | - | 0.00001913 | 0.00000611 | - | - |
| HLKn | - | <0.001 | - | 2.568 | 0.494 | - | - | 0.248 | 0.256 | - | - |
| HLKw | - | <0.001 | - | 0.00645 | 0.00231 | - | - | 0.00161 | 0.00244 | - | - |

\* variance component at least three times greater than its standard error
s.e.- standard error
<sub>x</sub>- ANOVA analysis only for infection condition

Correlations (based on the mean values for lines over locations) between the studied traits and FHBi were statistically significant in two types of treatments but values were generally low (Table 3). FHBi was negatively correlated with NSS, NGS, Sterility, Density, GWS, GY, HD and LSt at least in one type of treatment. Significant positive correlations were recorded between FHBi and Sterility in both control and infected conditions. Correlations between FHBi and *Fusarium* severity parameters (FDKn, FDKw, HLKn, HLKw) were significant under both type of treatments (an exception: HLKw in control condition). Positive, marginal correlations were found between FHBi and traits linked to FDK (FDKw and FDKn). HLKn and HLKw showed moderate correlations with FHBi.

**Table 3.** Correlation coefficients between FHB and studied traits recorded in two types of treatments.

| Trait | Treatment |  |
| --- | --- | --- |
|  | Infection | Control |
| NSS | -0.46 | -0.32 |
| NGS | -0.52 | -0.35 |
| LS | n.s. | n.s. |
| Sterility | 0.52 | 0.31 |
| Density | -0.43 | -0.27 |
| GWS | -0.44 | -0.35 |
| GY | -0.25 | -0.32 |
| TGW | n.s. | n.s. |
| Stature | -0.26 | n.s. |
| HD | -0.46 | -0.42 |
| LSt | -0.28 | n.s. |
| DON | n.s. | n.s. |
| FDK<sub>n</sub> | 0.25 | 0.26 |
| FDK<sub>w</sub> | 0.28 | 0.23 |
| HLK<sub>n</sub> | -0.46 | -0.42 |
| HLK<sub>w</sub> | -0.48 | n.s. |

n.s.- not significant
Correlations shown are significant at the $P < 0.01$ level

### Linkage map construction

The constructed genetic map comprised of 1947 SNPs distributed on seven linkage groups. The map length was 1678 cM with an average marker interval of 0.86 cM. The shortest chromosome was 6H, which harbored 250 markers with a genetic length of 141 cM and an average interloci distance of 0.56 cM. The longest chromosome was 2H, and it harbored 368 markers with a genetic length of 291 cM and an average interloci distance of 0.79 cM. The number of markers, marker density and map length for individual chromosomes are listed in Table 4.

**Table 4.** Map details across each chromosome.

|  | Chromosome |  |  |  |  |  |  | Total |
| --- | --- | --- | --- | --- | --- | --- | --- | --- |
|  | 1H | 2H | 3H | 4H | 5H | 6H | 7H |  |
| Number of mapped markers | 156 | 368 | 324 | 329 | 324 | 250 | 196 | 1947 |
| Map length (cM) | 232 | 291 | 241 | 215 | 295 | 141 | 263 | 1678 |
| Mean distance between markers (cM) | 1.48 | 0.79 | 0.74 | 0.65 | 0.91 | 0.56 | 1.30 | 0.86 |

## QTL analysis

A total of 70 QTLs for all studied traits were found for the LCam population. The numbers of QTLs were 7, 24, 5, 6, 17, 4, and 7 for chromosomes 1H, 2H, 3H, 4H, 5H, 6H and 7H, respectively. Moreover, 46 QTLs presented main effects and 38 presented QTL × E interaction. The largest number of QTLs was detected for NSS and TGW (eight QTLs were identified for each trait), and the smallest for FDK (two QTLs were detected for FDKn and FDKw). 14 QTLs were classified as major loci and 56 QTLs were decribed as minor loci. Detailed information, including location, peak marker, additive effects and explained phenotypic variance for each QTL and trait is presented in S3 Table.

S3 Table. QTLs identified in the LCam population for the observed traits.

### Spike characteristics

For the number of spikelets per spike eight QTLs were detected, in chromosomes 1H, 2H, 3H, and 5H. The major QTL (QNSS.IPG-2H_1) on chromosome 2H (SNP marker BK_12) showed the most significant effect for this trait and explained a large proportion of the phenotypic variance (4.79-71.81%). In this case significant QTL × E interaction was noted. A second locus positioned at 98.67 cM on chromosome 5H also showed a highly significant association with NSS (LogP statistics = 16.95). Chromosome 1H was the location of the last major QTL (QNSS.IPG-1H_1) in the vicinity of marker BOPA1_4625-1413. The remaining five NSS QTLs showed minor effects. Out of the eight QTLs detected for NSS, three (QNSS.IPG-1H_2, QNSS.IPG-2H_2 and QNSS.IPG-3H_2) were associated with a significant increase in this trait contributed by Syrian parent alleles.

Five QTLs were found for the number of grains per spike. QNGS.IPG-2H was found in the vicinity of marker BK_12. This locus was positioned at 22 cM on chromosome 2H. The second major QTL was detected on chromosome 5H in the vicinity of marker BOPA2_12_30929. For this QTL no significant additive effects were recorded in NAD location. The other QTLs (QNGS.IPG-1H_1, QNGS.IPG-1H_2 and QNGS.IPG-5H_2) were classified as minor QTLs. All studied QTLs for NGS were with alleles of European genotype contributing to an increasing of number of grains per spike.

Three QTLs were reported for the length of spike (QLS.IPG-1H, QLS.IPG-2H and QLS.IPG-5H). All detected loci were classiefied as major (≥12.32% PVE) and the effects of these QTLs were stable over environments (treatments). All QTLs were associated wih a significant increase in LS contributed by Lubuski. The main QTL was found n chromosome 2H in the vicinity of marker BK_13. In total, five QTLs were identified for sterility. On chromosome 2H, two major QTLs were detected (QSte.IPG-2H_1 and QSte.IPG-2H_2). The first QTL, QSte.IPG-2H_1, was located in the vicinity of marker SCRI_RS_154030 and showed the highest LogP value of all detected QTLs controlling this trait. The second sterility QTL was located on chromosome 2H 5.6 cM from marker SCRI_RS_230497. One major QTL (QSte.IPG-5H_2) was detected on chromosome 5H. None of the mentioned QTLs had significant additive effects in control condition in LES location and in infection conditions in NAD location. On chromosome 7H, a minor QTL for sterility was identified − QSte. IPG-7H. All QTLs detected for this character were with alleles of Syrian genotype contributing to the increase in sterility with the exception of QSte.IPG-5H_1, where Lubuski alleles determined the increase. Interaction with the environment was found for all but one detected QTLs (an exeption was QSte.IPG-7H).

Six QTLs controlling density were detected on chromosomes 2H and 5H with a PVE ranging from 0.01 to 31.43%. Half of those QTLs displayed significant QTL × E interaction. The main QTL (LogP=15.17) was found on the upper arm of chromosome 2H mapped in marker BK_22. Concurrently, this QTL was the only locus associated with Density, where Lubuski alleles conferred a positive effect in increasing this trait, while the Syrian parent alleles at the other five QTLs contributed positively to Density. The second major QTL (QDen.IPG-2H_2) was also found on chromosome 2H at position 113.9 cM. On chromosome 2H two other minor QTLs were identified for Density QTL (QDen.IPG-2H_2 and QDen.IPG-2H_4) with a stable effect, mapped in the vicinity of BOPA1_5537-283. QDen.IPG-5H_1 was also found on chromosome 5H at position 93.9 cM. The additive effects of this QTL was signifiant only in two location (NAD and TUL). For Density two minor QTLs were found − QDen.IPG-2H_3 and QDen.IPG-5H_2, for which the smallest LogP values were recorded for Density in this study.

### Grain traits

Grain weight per spike was mapped to seven loci. The main GWS QTL (QGWS.IPG-2H_1) was found on chromosome 2H in the vicinity of marker BK_22. This locus, with a PVE ranging from 31.74 – 57.92%, was the only GWS QTL where no significant QTL × E interaction was detected. QGWS.IPG-5H was found on chromosome 5H and the nearest marker (BOPA1_4795-782) was 1.37 cM away from the corresponding QTL peak. Two other major QTLs (QGWS.IPG-7H_1 and QGWS.IPG-7H_2) controlling GWS were reported on chromosome 7H. Both of these QTLs had a significant additive effect only in single location. Minor QGWS.IPG-4H_2 was found on chromosome 4H at position 127.40 cM. The European parent contributed to the increase in GWS for all detected QTLs for this trait (exceptions: two QTLs – minor QGWS.IPG-2H_2 and major QGWS.IPG-4H_1 identified on chromosomes 2H and 4H, respectively).

Out of four QTLs found for grain yield, only one was classiefied as major (PVE>12.32%). In additon, no significant additive effects were noticed for any detected loci in infection conditions for NAD location. The main QGY.IPG-2H was located on chromosome 2H and linked to marker BK_22. No QTL × E interaction was found for GWS QTLs detected in the mapping population and in all cases positive alleles were attributed to European parents.

Eight QTLs were reported for thousand grain weight. QTGW.IPG-2H_1 and QTGW.IPG-4H_1 were identified on chromosomes 2H and 4H, respectively, but their additive effects were significant only in infection (LES) and control conditions (NAD). On chromosome 4H, TGW QTL was found with a stable and positive effect from the Lubuski genotype. QTGW.IPG-6H_2 locus on chromosome 6H was determined by Syrian parent genotype alleles contributing positively to TGW. In this locus no significant QTL × environment interaction for TGW was also observed. Major QTGW.IPG-7H_1 with stable effects from the CamB allele significantly increasing TGW was identified on chromosome 7H. On the same chromosome was found QTGW.IPG-7H_2, but the additive effects of this QTL were significant only in three treatments. QTGW.IPG-2H_2 and QTGW.IPG-6H_1, detected on chromosome 2H and 6H, respectively, were classiefied as minor QTLs.

### Heading day and height

Two QTLs (QHD.IPG-2H and QHD.IPG-5H) were reported for heading date. The main QTL was located on chromosome 2H in the vicinity of marker BK_22. The „late” allele (high HD value) was contributed by European parent. In contrast, at the second locus, classified as a minor QTL, the CamB alleles conferred a positive effect by increasing this trait. For both loci, no QTL × E interaction was detected.

Seven loci for length of main stem were found in the LCam population. The main locus (QLSt.IPG-2H_1) was detected on chromosome 2H in the vicinity of marker BK_13 at position 21 cM. This QTL explained a large portion of the variance for LSt (from 13.88 to 41.68%). The Lubuski alleles contributed to the increase in LSt at this locus. The second major QTL was reported on chromosome 1H with stable and positive effects on the length of the main stem contributed by the European parent genotype. QLSt.IPG-4H_2 and QLSt.IPG-5H were identified on chromosomes 4H and 5H, respectively. These QTLs were classified as major loci, but their additive effects were not significant in some treatments (e.g. control conditions in NAD location). Three minor LSt loci were found − QLSt.IPG-2H_2, QLSt.IPG-3H and QLSt.IPG-4H_1 detected on chromosomes 2H, 3H and 4H, respectively.

### Fusarium symptoms and DON content

Six QTLs were reported for the FHB index. The main QTL (QFHBi.IPG-2H_1) was found on chromosome 2H in the vicinity of marker BOPA1_5880-2547 at position 23.10 cM. The CamB alleles positively contributed to the increase in the FHB index at this locus and significant QTL × E interaction was detected for QFHBi.IPG-2H_1. On the same chromosome another FHBi QTL was reported which was located at position 87.70 cM but additive effects of this locus were significant only in one location (TUL). The next major locus (QFHBi.IPG-2H_3) was also detected on chromosome 2H with stable and positive effects of Syrian parent alleles responsible for increasing the FHBi. In contrast, the European parent alleles conferred a positive effect in increasing the FHBi at the locus found on chromosome 5H (QFHBi.IPG-5H). For this QTL no significant additive effects were detected in LES location. Two minor loci − QFHBi.IPG-3H and QFHBi.IPG-7H were reported on chromosome 3H and 7H, respectively.

Four QTLs were found for traits linked to *Fusarium* damaged kernels. These loci were located on chromosomes 5H and 6H. The main QFDKn.IPG-5H was detected in the vicinity of SCRI_RS_165578, where Lubuski genotype significantly increased the FDKn. The second major locus (QFDKw.IPG-5H) was identified at the position 87.80 cM and showed positive effects on this trait contributed by European parent alleles. Two remaining loci found on chromosome 6H (QFDKn.IPG-6H and QFDKw.IPG-6H) were classified as minor QTLs.

Five QTLs were detected for traits associated with healthy looking kernels (HLKw and HLKn). The main QHLKn.IPG-2H_2 was found on the short arm of chromosome 2H (marker BK_13) and showed stable and positive effects of Lubuski genotype alleles which contributed to the increase in HLKn. Two minor QTLs were recorded for HLKn on chromosomes 2H and 5H. For both loci no significant QTL x E interaction was detected. Two major loci (QHLKw.IPG-2H and Q_HLKw.IPG-7H) were found for the trait HLKw. The Lubuski alleles were responsible for increasing HLKw in both loci but only one QTL (QHLKw.IPG-2H) had stable effects.

### Co-localized or pleiotropic QTLs

A total of eight chromosomal regions (named A-G) harboring QTLs for studied traits were assigned. These regions (hotspots), listed in Table 5, were designed based on their proximity to each other (0-5 cM).

**Table 5.** Regions harboring QTLs for studied traits with the names of the nearest SNP markers.

| name of hotspo | Trait | QTL ID | Chromosome | Position (cM) | Nearest marker |
| --- | --- | --- | --- | --- | --- |
| A | NSS | QNSS.IPG-1H_1 | 1H | 0,00 | BOPA1_4625-1413 |
|  | NGS | QNGS.IPG-1H_1 | 1H | 0,00 | BOPA1_4625-1413 |
|  | LS | QLS.IPG-1H_1 | 1H | 0,00 | BOPA1_4625-1413 |
|  | LSt | QLSt.IPG-1H_1 | 1H | 0,00 | BOPA1_4625-1413 |
|  | GY | QGY.IPG-1H_1 | 1H | 0,00 | BOPA1_4625-1413 |
| B | LS | QLS.IPG-2H_1 | 2H | 21,00 | BK_13 |
|  | LSt | QLSt.IPG-2H_1 | 2H | 21,00 | BK_13 |
|  | NSS | QNSS.IPG-2H_1 | 2H | 22,00 | BK_12 |
|  | NGS | QNGS.IPG-2H_1 | 2H | 22,00 | BK_12 |
|  | Density | QDen.IPG-2H_1 | 2H | 22,00 | BK_12 |
|  | GWS | QGWS.IPG-2H_1 | 2H | 22,00 | BK_12 |
|  | GY | QGY.IPG-2H_1 | 2H | 22,00 | BK_12 |
|  | HD | QHD.IPG-2H_1 | 2H | 22,00 | BK_12 |
|  | HLKw | QHLKw.IPG-2H_1 | 2H | 22,00 | BK_12 |
|  | FHBi | QFHB.IPG-2H_1 | 2H | 23,10 | BOPA1_5880-2547 |
| C | Density | QDen.IPG-2H_2 | 2H | 225,26 | BOPA2_12_10937 |
|  | LSt | QLSt.IPG-2H_2 | 2H | 225,26 | BOPA2_12_10937 |
|  | GWS | QGWS.IPG-2H_2 | 2H | 228,70 | BOPA2_12_10937 |
|  | TGW | QTGW.IPG-2H_2 | 2H | 228,70 | BOPA2_12_10937 |
|  | NSS | QNSS.IPG-2H_2 | 2H | 229,80 | SCRI_RS_174051 |
| D | TGW | QTGW.IPG-4H_1 | 4H | 127,40 | BOPA1_2196-195 |
|  | GWS | QGWS.IPG-4H_1 | 4H | 127,40 | BOPA1_2196-195 |
| E | Density | QDen.IPG-5H_1 | 5H | 93,90 | SCRI_RS_184066 |
|  | FHBi | QFHB.IPG-5H_1 | 5H | 95,60 | SCRI_RS_184066 |
|  | NGS | QNGS.IPG-5H_1 | 5H | 97,30 | BOPA2_12_30929 |
|  | LSt | QLSt.IPG-5H_1 | 5H | 97,30 | BOPA2_12_30929 |
|  | LS | QLS.IPG-5H_1 | 5H | 97,30 | BOPA2_12_30929 |
|  | NSS | QNSS.IPG-5H_1 | 5H | 98,67 | SCRI_RS_235055 |
| F | GY | QGY.IPG-5H_2 | 5H | 285,20 | BOPA2_12_30533 |
|  | NSS | QNSS.IPG-5H_2 | 5H | 286,90 | SCRI_RS_206867 |
|  | NGS | QNGS.IPG-5H_2 | 5H | 286,90 | SCRI_RS_206867 |
|  | HLKn | QHLKn.IPG-5H_2 | 5H | 288,00 | SCRI_RS_165919 |
| G | GWS | QGWS.IPG-7H_1 | 7H | 119,80 | SCRI_RS_159555 |
|  | TGW | QTGW.IPG-7H_1 | 7H | 119,80 | SCRI_RS_159555 |

Five QTLs were reported in region A located on the top of chromosome 1H, namely associated with SNP BOPA1_4625-1413. Region B identified on the upper arm of chromosome 2H contained to 10 loci. In most cases, QTLs from region B were detected in the vicinity of marker BK_12 (Fig 4). Out of the five QTLs detected in region C assigned on the same chromosome, four were found in the vicinity of marker BOPA2_12_10937. Region D harbored two QTLs found at the same position (127.40 cM) but these loci were linked to different SNP markers. Region E (chromosome 5H) harbored six QTLs – both QDen.IPG-5H_1 and QFHB.IPG-5H were found in this region in the vicinity of marker SCRI_RS_184066 and both QNGS.IPG-5H_1 and QLSt.IPG-5H were detected in the vicinity of marker BOPA2_12_30929. On the same chromosome, the next region was noted (named region F). Out of the four QTLs reported on this region, two were found in the vicinity of marker SCRI_RS_206867. Region F on chromosome 7H harbored two loci associated with marker SCRI_RS_159555.

**Fig 4.**
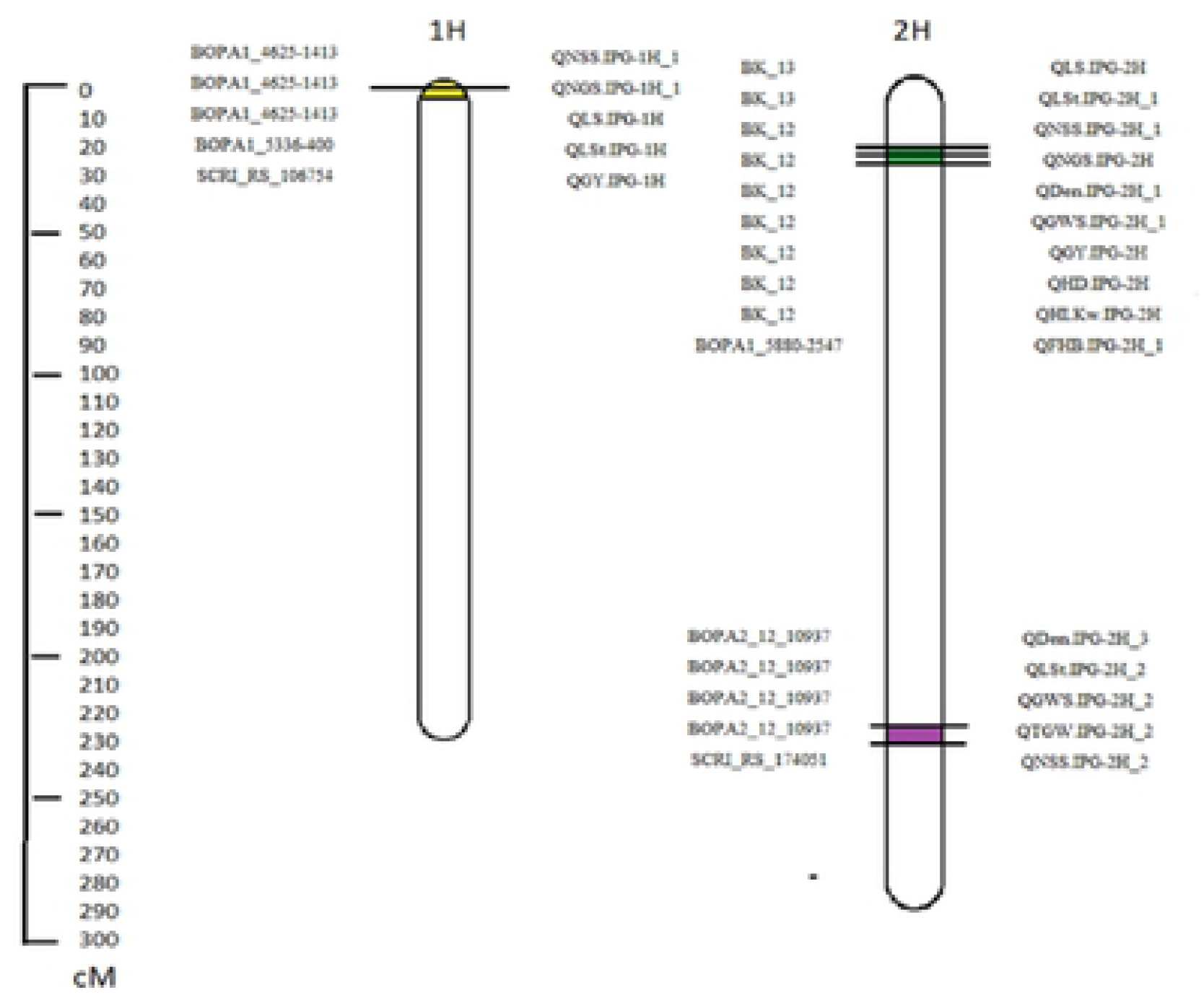
Locations of regions (A – chromosome 1H and B,C – chromosome 2H) harboring QTLs detected for studied traits in the LCam population. Ppd-H1 gene localisation and corresponding markers were shown. Genetic distance scale in centiMorgan (cM) is place in the left margin.

An overrepresentation analysis was performed to identify enriched Gene Ontology (GO) terms – cellular component, molecular function and biological process - associated with regions listed in Table 5, containing QTLs connected to FHB. There are multiple genes that are in the regions - hotspot associated with traits analyzed in this study. Based on gene annotation and literature studies, selected candidate genes related to disease resistance were shown in Table 6.

**Table 6.**
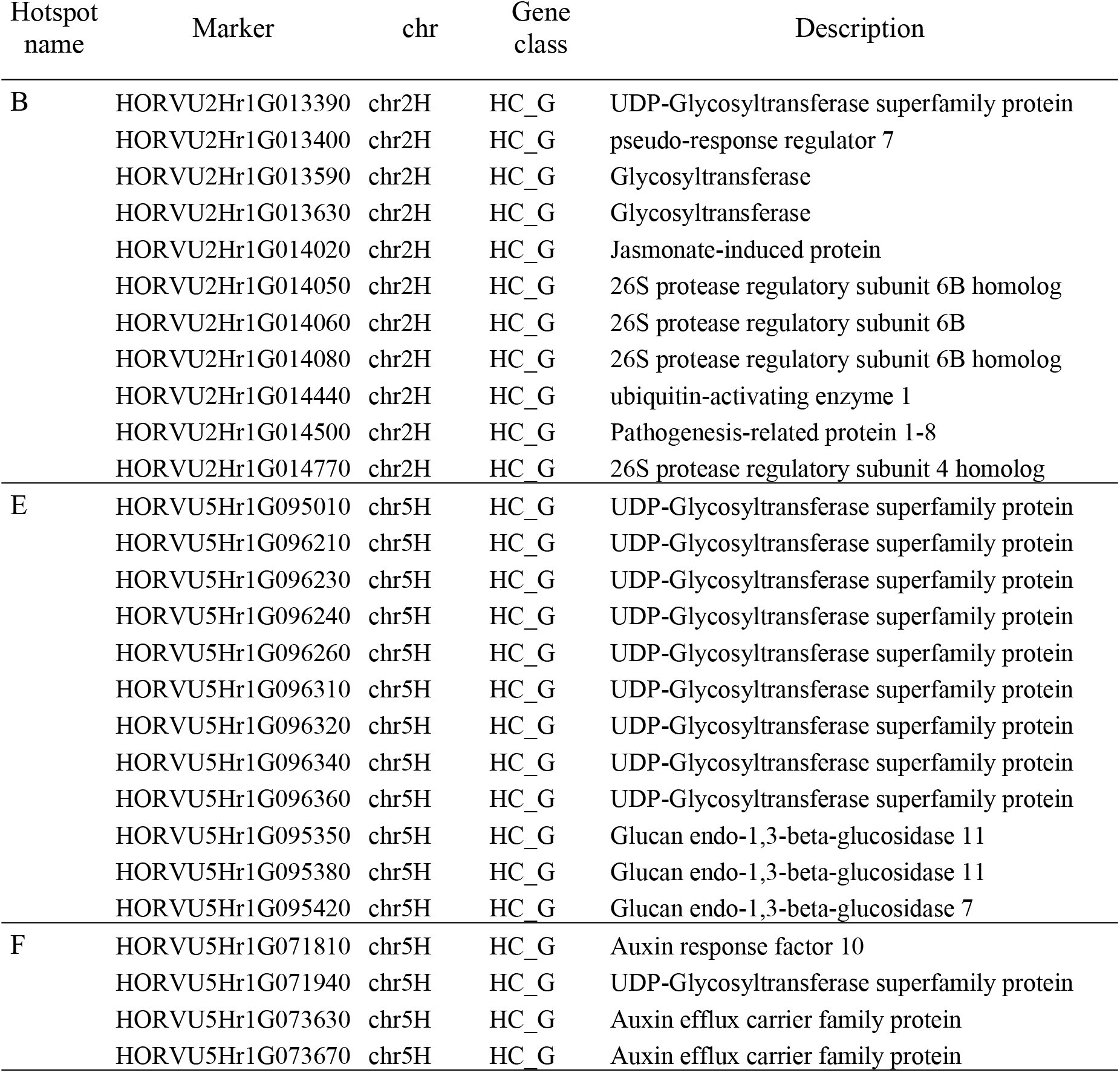
Selected candidate genes identified by using the Barleymap database (http://floresta.eead.csic.es/barleymap).

| Hotspot name | Marker | chr | Gene class | Description |
| --- | --- | --- | --- | --- |
| B | HORVU2Hr1G013390 | chr2H | HC_G | UDP-Glycosyltransferase superfamily protein |
|  | HORVU2Hr1G013400 | chr2H | HC_G | pseudo-response regulator 7 |
|  | HORVU2Hr1G013590 | chr2H | HC_G | Glycosyltransferase |
|  | HORVU2Hr1G013630 | chr2H | HC_G | Glycosyltransferase |
|  | HORVU2Hr1G014020 | chr2H | HC_G | Jasmonate-induced protein |
|  | HORVU2Hr1G014050 | chr2H | HC_G | 26S protease regulatory subunit 6B homolog |
|  | HORVU2Hr1G014060 | chr2H | HC_G | 26S protease regulatory subunit 6B |
|  | HORVU2Hr1G014080 | chr2H | HC_G | 26S protease regulatory subunit 6B homolog |
|  | HORVU2Hr1G014440 | chr2H | HC_G | ubiquitin-activating enzyme 1 |
|  | HORVU2Hr1G014500 | chr2H | HC_G | Pathogenesis-related protein 1-8 |
|  | HORVU2Hr1G014770 | chr2H | HC_G | 26S protease regulatory subunit 4 homolog |
| E | HORVU5Hr1G095010 | chr5H | HC_G | UDP-Glycosyltransferase superfamily protein |
|  | HORVU5Hr1G096210 | chr5H | HC_G | UDP-Glycosyltransferase superfamily protein |
|  | HORVU5Hr1G096230 | chr5H | HC_G | UDP-Glycosyltransferase superfamily protein |
|  | HORVU5Hr1G096240 | chr5H | HC_G | UDP-Glycosyltransferase superfamily protein |
|  | HORVU5Hr1G096260 | chr5H | HC_G | UDP-Glycosyltransferase superfamily protein |
|  | HORVU5Hr1G096310 | chr5H | HC_G | UDP-Glycosyltransferase superfamily protein |
|  | HORVU5Hr1G096320 | chr5H | HC_G | UDP-Glycosyltransferase superfamily protein |
|  | HORVU5Hr1G096340 | chr5H | HC_G | UDP-Glycosyltransferase superfamily protein |
|  | HORVU5Hr1G096360 | chr5H | HC_G | UDP-Glycosyltransferase superfamily protein |
|  | HORVU5Hr1G095350 | chr5H | HC_G | Glucan endo-1,3-beta-glucosidase 11 |
|  | HORVU5Hr1G095380 | chr5H | HC_G | Glucan endo-1,3-beta-glucosidase 11 |
|  | HORVU5Hr1G095420 | chr5H | HC_G | Glucan endo-1,3-beta-glucosidase 7 |
| F | HORVU5Hr1G071810 | chr5H | HC_G | Auxin response factor 10 |
|  | HORVU5Hr1G071940 | chr5H | HC_G | UDP-Glycosyltransferase superfamily protein |
|  | HORVU5Hr1G073630 | chr5H | HC_G | Auxin efflux carrier family protein |
|  | HORVU5Hr1G073670 | chr5H | HC_G | Auxin efflux carrier family protein |

## Discussion

It is widely known that a mapping population derived from parents divergent in genetic composition allows high performance QTL analysis. In this study, RILs (named LCam) derived from a European variety and a Syrian breeding line were used for QTL analysis. Both parental forms were differentiated in terms of stature, grain yield, HD and resistance/tolerance to biotic and abiotic stresses [37, 38]. CamB is unadapted to the Central Europe region and has undesired agonomical trats such as early heading and tall stature. Lubuski is an old cultivar with agro-morphological-physiological characters adapted to Polish climatic conditions during a long cultivation period. With the aim of providing the genetic variability between the parents of the mapping populaion and increasing the chance to indentifying loci linked to FHB we conducted field experiments using RILs derived from a cross between Lubuski and CamB genotypes.

FHB, caused by *Fusarium culmorum*, is a very important disease of crops globally [9]. Damage caused by *Fusarium* species fungus includes reduced grain yield, and reduced grain functional quality, and results in the presence of the mycotoxin deoxynivalenol in FDK (and also in grains without any visible symptoms). The development of FHB resistant crop cultivars is an important component of integrated breeding management [54, 55]. The objective of this investigation was to identify QTLs for traits linked to yield performance in a recombinant inbred line population grown under a disease-free environment and under *Fusarium* infection conditions.

FHB infection can be evaluated in different ways. In field conditions, FHB can be determited, among others, by visual inspection of the percentage of infected spikelets [56]. This percentages can be used to determine an FHB index [57]. In this study, different tools were employed to estimate disease severity: FHBi was calculated as: (spike infection in % percentage of spikelets affected within a spike) / (percentage of infected spikes per plot). After harvest, percentage both FDK and HLK as described by the visual symptom score of the kernels and the weight of the kernels were evaluated. In addition, DON concentration was quantified.

In this study Lubuski was less susceptible to FHB than Syrian parent in all type of conditions in terms of DON accumulation. On the other hand, we observed a higher FHBi value for Lubuski plants in infection conditions. DON tests of grains harvested from the LCam population showed that some RILs showed lower DON content values than CamB, while some other RILs were more susceptible than Lubuski. In all types of conditions, the mean values of agronomic traits performed as expected – biotic stress conditions impaired yields. In control conditions, DON contamination represents the natural occurrence of FHB [58] and the level of mycotoxin accumulation varied significantly from the ones observed in LCam plants grown in infection conditions.

Most of the correlation coefficients among the FHBi and other studied characters were negative an statistically different from zero (P<0.01). The Pearson correlation coefficient was also significantly negative between FHBi and the two main traits of our interest – HD and LSt, which is in agreement with previous studies [59–61], where plants with lower FHB severities have usually been characterized by late heading and tall stature. Late-maturing plants may head during a time in the summer less suitable for infection, and tall plants avoid higher concentrations of inoculum near the surface of the soil [62]. Another study, conducted by Mesfin et al. [24] concluded that late HD may be linked to FHB resistance since the heads experience less exposure time to fungal spores.

Visual ratings for FHB in barley plants are usually conducted just before the spikes begin to lose chlorophyll so disease symptoms can be easily counted. In some years, there are favorable conditions for *Fusarium* growth and DON accumulation throughout plant senescence, reducing correlations between FHB and DON because the FHB score does not accurately reflect the final disease level [63]. It is well known that symptomless spikes can be contaminated with DON [64]. Traditionally, mycotoxin determination has mainly been performed by chromatographic techniques [65, 66]. ELISA has been proposed as an alternative method to visual scoring and DON quantification for mesuring FHB [67] (Hill et al. 2008). Expression of DON by fungal myceium is environmentally dependent [68], whereas the expression of monoclonal antibody-specific mycelial proteins is not [69]. In consequence, ELISA and DON values are not always closely correlated. In our study, no correlation was found between FHBi and DON level content, which can be explained by this phenomenon.

Agronomic traits related to spike characters (e.g., spike density and sterility) have been reported to be linked to FHB resistance but the association between the traits and FHB vulnerability seems to be unclear. Steffenson et al. [70] reported that FHB severity was apparently higher in dense spike NILs than in lax spikes. A negative correlation between FHB severity and spike density has been recorded in an experiment conducted on a population derived from two-row and six-row barley plants [71]. In contrast, in the study conducted by Yoshida et al. [72] on barley NILs spike density had little or no effect. Ma et al. [73] also reported an association between lax spike and FHB reaction. Lax spikes may be related to FHB resistance due to their specific architecture that retains, presumably, less moisture within the whole spike. This decreases the pace of fungus spread [4]. In the presented study negative correlations were detected between the traits Density and FHBi, indicating that spike compactness may be one of the factors enhancing FHB susceptibility. A positive correlation was recorded between Sterility and FHBi for LCam plants, which means that FHB infection has negative effects on seed development, as expected.

Single nucleotide polymorphisms (SNPs) that are widely distributed throughout the genome have been used in QTL mapping [74, 75]. In our previous study using 1536 SNP markers, we developed genetic map which allowed us to identify a set of QTLs linked to different agronomic traits (among others: HD and plant height) in field [38] and greenhouse conditions [37]. In this study, by using a 9K SNP array, we improved the QTL mapping resolution due to increased marker density.

The SNP markers were distributed across all seven linkage groups in the LCam mapping population. Marker order and distances for SNPs generally matched previously published barley maps [40, 76]. The genetic map consisting of 1947 SNPs which has been developed in this investigation, covering 1678 cM, is larger than other maps (e.g., constructed by Wang et al. [77] − 1375.8 cM.

Many bi-parental mapping studies have been conducted on barley to explain the genetic architecture of resistance to FHB and DON accumulation and to identify molecular markers that could be useful in breeding [24, 31, 78, 79]. FHB resistance has frequently been found to be associated with plant morphology parameters, especially plant height, spike architecture, anther extrusion and HD acting mainly as passive resistance factors. For this reason, the LCam population was also evaluated for HD, plant height, spike compactness and other traits, which seem to be important form an agronomic point of view. Numerous QTL mapping studies that have been conducted in different crop species have revealed that QTLs associated with FHB resistance are coincident with QTLs linked to various agronomic and morphological traits [4, 24, 79]. In our investigation, a set of 70 QTLs were detected in seven barley chromosomes. A higher number of QTLs for agronomic traits was found on chromosome 2H, where the greatest number of FHB-linked QTLs was also identified.

QTLs for FHB resistance have been found on all seven barley chromosomes [24, 31, 78– 81]. For most of the resistance varieties, QTLs associated with FHB were detected on the long arm of chromosome 2H [30, 31, 82]. In addition, QTLs for disease resistance and reduced DON concentration have been linked to spike morphology controlled by *vrs1* and a major HD locus (P*pd-H1*) in numerous studies [80]. The number of detected QTLs varies depending on the type of research, ranging from only one in the study conducted by Mesfin et al. [24] to two [4, 31, 73] or even to 10 QTLs [22]. For many FHB regions in the barley genome, QTLs for DON concentration have been detected both for barley [81, 82] and for wheat [83, 84] but this kind of coincidence has not been reported as significant in all studies [30]. Identification of QTLs linked to FHB symptoms has been confounded by agronomic characters such as HD, plant height or properties associated with spike morphology [24, 79]. Hence, mapping of traits characterized by strong phenotypic correlations constitutes a challenge in terms of pleiotropy/linkage. Massman et al. [80] have summarized previously described FHB regions and showed all detected QTLs in a graph associated with genome location (bin). The QTLs were located on chromosome 2H at three different spots (bin 8, bin 10 and bin 13-14). In our investigation, six QTLs related do FHBi were found. Out of these QTLs, three were identified on chromosome 2H at position 23.1, 87.7 and cM corresponding to previously mentioned bin locations. Three other loci: QFHB.IPG-3H, QFHB.IPG-5H and QFHB.IPG-7H were found on chromosomes 3H, 5H and 7H, respectively. QFHB.IPG-2H_1 was found on the short arm of chromosome 2H in the vicinity of SNP marker BOPA1_5880-2547 and this explains the largest percentage of phenotyping variance (3.69 – 30.69) of all detected FHB QTLs. The Syrian parent alleles positively contributed to the increase in FHBi at this locus, which is in accordance with previous studies, in which early heading plants were vulnerable to FHB symptoms. In our study, the main QTL for HD was located on chromosome 2H in the vicinity of marker BK_12 at position 22 cM, shifted 1.1 cM from marker BOPA1_5880-2547. According to Turner at al. [85] the most significant SNP marker (BK_12) is directly located within the *Ppd-H1* gene, which is the main determinant of response to long day conditions in barley. The 2Hb8 QTL is also considered a major locus for resistance to FHB and DON accumulation [86]. Delayed head emergence may increase the likelihood that the host will escape infection by the pathogen [71, 87]. On the other hand, late heading is undesirable in breeding programs addressed to arid regions [88]. Plants with lower FHB severities usually have one or more of the following traits: late heading, increased height and two-rowed spike morphology [59, 60, 70]. Although tall plants are usually more resistant to disease than short plant [73], heading date can be either negatively [71, 73] or positively correlated with DON content in the seeds [22, 24]. The main QTL associated with heading and located on chromosome 2H (Q.HD.LC-2H), was also identified at SNP marker 5880–2547, in our previous study [37]. SNP 5880–2547 was the closest marker to QTLs associated with plant architecture, spike morphology and grain yield in the mentioned experiment. None of the detected QTLs for FHBi in the current study were classified as major loci (the heighest value of LogP value was 8.14 recorded for QFHB.IPG-2H_3), but this could be explained by the fact that the identification of many minor QTLs associated with the model of complex traits that predicts an exponential decay of QTL effects only a few loci but has large effects [89].

Plant height is under polygenic control and represents one of the most important agronomic traits for barley [90, 91]. The right timing of flowering time allows optimal grain development with regard to the availability of heat, light and water, while semi-dwarf cereals allocate more resources into grain production tan taller plants and show reduced losses through lodging [92, 93]. In addition, due to increasing of moisture content of the plants, lodging causes the infection expansion [94]. In the current study were detected seven loci for LSt. The main locus (QLSt.IPG-2H_1) was detected on chromosome 2H in the vicinity of marker BK_13, which coincided with the main HD QTL. In this study only one locus was found on chromosome 3H, where *sdw1/denso* has been located in our previous investigations [38, 90, 95]. There is a gradient at ascospore concentration from the soil surface to upper part of plant stem. Thus, short plant tend to have higher FHB infection level [96], which is in accordance with our results.

In barley, spike length and spike characters like number of grains and spikelets per spike are perceived as an important agromorphological traits due a direct impact on crop yield [97]. The spike architecture has significant influence on yield and might alter the spike microenvironment making it less favorable for fungal infection [98]. In the current study six QTLs linked to Density were found. Out of six detected QTLs, four loci were found on chromosome 2H. The major QTL (QDen.IPG-2H-1) was located on the short arm of 2H in the vicinity of marker BK_12. Two QTLs related to the density of the spike were found on chromosome 5H. In most cases CamB alleles contributed positively to this trait. In many studies plants with lax spikes have been reported as less vulnerable for fungal infection [82, 98]. On the other hand, Yoshida et al. [62] found no differences between genotypes when compared barleys with normal and dense type of spike. Steffenson et al. [70] showed that FHB severity was higher in dense spike NILs than in lax spike plants but no significant differences were found. Langevin et al. [99], conducting the study using barley with two- and six-row type of spike, concluded that the high level of DON contamination observed among dense spikes accured mainly because of direct contact of the florets. To summarize, results for the association between disease severity and spike architecture of the barley plants are not consistent.

The genes encoding UDP-Glycosyltransferase superfamily protein were found for all hotspots containing QTLs linked to FHB (FHBi, HLKn, HLKw). Plant uridine diphosphate (UDP)-glucosyltransferases (UGT) catalyze the glucosylation of xenobiotic, endogenous substrates and phytotoxic agents produced by pathogens such as mycotoxins [100, 101]. The studies have shown that plant UDP-glucosyltransferase genes have significant role in plant resistance both to biotic and abiotic stresses [102, 103]. Poppenberger et al. [104] demonstrated that DON resistance can be achieved by the enzymatic conversation (a natural detoxification process in plants called glycosylation) of the toxin into the non-toxic form (DON-3-0-glucoside) by UDP-glucosyltransferase. It is also worth to mention that in our study 10 records have been annotated for region E, where FHBi QTL was found on chromosome 5H. Recently the HvUGT-10 W1 gene has been isolated from an FHB resistant barley variety conferred FHB tolerance [102].

Region B, as tightly linked to the QTLs identified for FHBi and HD in this study, has been annotated also as pathogenesis-related protein 1-8 (HORVU2Hr1G014500). These type of proteins belong to the antimicrobial compounds playing an important role in defense response against fungal infection. Induction of PR (Pathogen-related)-proteins has been found in many plant species belonging to various families [105]. For instance, Pritsch et al. [106] observed that the transcripts of defense response genes, peroxidase and PR-1 to −5, accumulated as early as 6 to 12 h after wheat spikes were inoculated with *F. graminearum*.

Another gene annotation linked to disease resistance was shown for region B. Glucan endo-1,3-beta-glucosidase may provide a degree of protection against microbial invasion of germinated barley grain through its ability to degrade fungal cell wall polysaccharides, according to Balasubramanian et al. [107].

Data obtained from Barley Map Floresta database allowed us to annotate both in region F and B candidate genes related to two types of phytohormones, which play pivotal roles in regulation of this defence network [108]. Several regions identified in our analysis included annotations associated with auxin (HORVU5Hr1G071810 - auxin response factor 10; HORVU5Hr1G073630, chr5H, HORVU5Hr1G073670 - auxin efflux carrier family protein) in region F. The roles of auxins in plant-pathogen interactions have also been described in recent years [109, 110]. Another phytohormone annotation (jasmonate-induced protein) was present in the region B. Jasmonic acid (JA) is considered to be a critical phytohormone for plant defense response against pathogens [111]. For instance, Gottwald et al. [112] suggested that Jasmonate and ethylene dependent defense and suppression of fungal virulence factors are major mechanisms of FHB resistance in wheat.

In the region B were found also following annotations:pseudo-response regulator 7 (HORVU2Hr1G013400), 26S protease regulatory subunit 6B (HORVU2Hr1G014050, HORVU2Hr1G014060 and HORVU2Hr1G014080) and ubiquitin-activating enzyme 1 (HORVU2Hr1G014440). All these annotations are linked to *HvPpd-H1*, which provides adaptation to photoperiod in barley. The barley Ppd-H1 gene is homologous to Arabidopsis PRR3/PRR7 and mediates the acceleration of development in long-days [113]. The Ppd-H1/PRR37 allele is the major determinant of photoperiod response in barley and is the putative AtPRR7 orthologue [85]. Proteasomes are involved in the degradation of ubiquitin-tagged proteins. Alterations in proteasome subunits were found in several proteomic studies dealing with both abiotic and biotic stresses [114, 115].

Ubiquitin is well established as the major modifier of signaling in eukaryotes. The main characteristic of ubiquitination is the conjugation of ubiquitin onto lysine residues of acceptor proteins [116]. In most cases, the targeted protein is degraded by the 26S proteasome, the major proteolysis machinery in eukaryotic cells. The ubiquitin– proteasome system is responsible for removing most abnormal peptides and short-lived cellular regulators. This allows cells to respond rapidly to intracellular signals and changing environmental conditions. In our study, annotations linked to 26S protease regulatory subunits were found in region B (HORVU2Hr1G014050, HORVU2Hr1G014060, HORVU2Hr1G014080 and HORVU2Hr1G014770).

## Conclusions

Results from the research conducted recently have revealed that most of the resistant barley genotypes showed QTL linked to FHB on the long arm of 2H chromosome, the first a coincident QTL for HD and the second associated with *vrs1* gene. In our study, two major QTLs (QFHB.IPG-2H_1 and QFHB.IPG-2H_2) found on chromosome 2H were located in the similar positions as the loci detected in previus studies (30, 86). A major confounding problem in mapping loci for FHB resistance is that QTLs for the agromorphological traits (HD, length of stem and spike type) are often coincident with the loci for disease resistance and they may interfere with resistance evaluation [82, 83]. Furthermore, it can be difficult to reveal the genetic architecture of these traits (linked QTL or pleiotropy) when many QTL are identified at the same locus. On the other hand, our results support a major assumption that plant architecture and inflorescence traits are associated significantly with FHB severity.

Out of six detected FHB QTLs in the current study, four were not classified as a loci localisated in the hotspots, where many yield-related loci were detected. Although, in previously conducted studies QTLs associated with FHB were found on chromosomes 3H, 5H and 7H, QTLs identified in our investigation appear to be unique for FHB symptoms. Thus, the barley genotypes carrying these QTLs may be used in breeding programs without the confounding effects from another yield-related traits.

## Supporting Information

S1 Table. The mean values for studied traits for parental cultivars. (DOC)

S2 Table. The mean values for studied traits for RILs. (DOC)

S3 Table. QTLs identified in the LCam population for the observed traits. (DOC)

## Acknowledgments

This work was financially supported by Polish Ministry of Agriculture and Rural Development (grant no HOR hn-501-19/15 Task 88).

## Author Contributions

PO, AK, KM conception and design of the study; AK funding acquisition, project coordination; PO, MK, MR, DJ plant breeding and stresses application; MK, PO, KM, AK sampling of plant material; PK, HĆK methodology validation, statistics and computations; PO, AK, KM, MK, DJ samples preparation, laboratory work and analyses; PO, AK, KM, TA, MS, PK data analysis and interpretation, manuscript writing, revision and editing; PO, AK, KM, MK, TA, MS, PK, HĆK, MR and DJ contributed to the final version of the manuscript.

